# Bayesian Aggregation of Multiple Annotations Enhances Rare Variant Association Testing

**DOI:** 10.1101/2025.03.02.641062

**Authors:** Antonio Nappi, Liubov Shilova, Theofanis Karaletsos, Na Cai, Francesco Paolo Casale

## Abstract

Gene-level rare variant association tests (RVATs) are essential for uncovering disease mechanisms and identifying potential drug targets. Advances in sequence-based machine learning have expanded the availability of variant pathogenicity scores, offering new opportunities to improve variant prioritization for RVATs. However, existing methods often rely on rigid models or analyze single annotations in isolation, limiting their ability to leverage these advances. To address this, we introduce BayesRVAT, a Bayesian framework for RVATs that models variant effects using multiple annotations. By specifying priors on variant effects and estimating gene-trait-specific burden scores through variational inference, BayesRVAT flexibly accommodates diverse genetic architectures. In simulations, BayesRVAT improved power while maintaining statistical calibration. When applied to real data from the UK Biobank, BayesRVAT detected 10.2% more associations with blood traits than the next-best method and uncovered novel gene-disease associations in analyses of eight disease traits, including a link between *PRPH2* and retinal disease. These results highlight BayesRVAT’s potential to enhance rare variant discovery and improve gene-trait association studies.

## Introduction

Understanding the role of rare genetic variants is crucial for uncovering disease mechanisms and identifying potential therapeutic targets. Prioritizing variants with lower frequencies and potential functional impact, rare variant analyses tend to provide a more interpretable approach compared to common variant studies (Cirulli et al. 2020; McCaw et al. 2023).

Gene-based rare variant association studies (RVATs) aim to detect associations between rare variants in a gene and traits of interest. Traditionally, gene-based RVATs have been performed using burden tests, which aggregate likely deleterious variants into a gene burden score and then regress these scores against trait values across individuals in a formal gene-level association test (Backman et al. 2021; Jurgens et al. 2022; Karczewski et al. 2022; Madsen and Browning 2009; Li and Leal 2008; Brandes et al. 2020). Likely deleterious variants within a gene are typically identified based on consequence annotations (e.g., protein-truncating variants tend to have stronger effects than missense variants) (McLaren et al. 2016) and functional effect prediction scores such as PolyPhen-2 and SIFT (Adzhubei et al. 2010; Karczewski et al. 2022; Kumar et al. 2009). More recently, machine learning models trained on biological sequences to predict functional and structural properties have expanded the availability of variant pathogenicity prediction scores (Sundaram et al. 2018; Jaganathan et al. 2019; Wagner et al. 2023; Ghanbari and Ohler 2020; Zhou and Troyanskaya 2015; Brandes et al. 2023; Cheng et al. 2023), providing new opportunities for improved variant prioritization.

Recent burden test models can incorporate multiple variant annotations directly implementing the concept of an *allelic series*, where increased likelihoods of gene disruptions correspond to stronger phenotypic effects (McClintock 1944; Musunuru and Kathiresan 2019). For instance, COAST weights variants based on their predicted deleteriousness (McCaw et al. 2023), while DeepRVAT employs a data-driven approach to learn aggregation functions from multiple annotations using a neural network (Clarke et al. 2024). Despite these improvements, a unified framework that jointly models multiple annotations while allowing flexible, gene- and trait-specific aggregation is still missing.

To address this, we present BayesRVAT, a Bayesian RVAT framework that flexibly aggregates variant effects using multiple annotations. Inspired by the concept of allelic series, BayesRVAT models variant effects as a function of multiple annotations and aggregates them in a gene- and trait-specific manner through Bayesian inference, capturing how different annotations shape variant burden. To compute association P values within this Bayesian framework, we introduce an approximate likelihood ratio testing. We validate BayesRVAT through simulations and analyses of quantitative and binary traits from the UK Biobank, demonstrating improved performance over existing gene-level RVAT strategies.

## Results

### Bayesian Aggregation for Rare Variant Association Testing

Gene-level burden tests aggregate the effects of rare variants within a gene into a single burden score, which is then tested for association with the trait of interest. Formally, given trait values *y* ∈ *R* ^*N*^ for *N* individuals, genotype matrix *X* = [*x*_1_, …, *x*_*N*_] ^*T*^∈ *R* for *S* rare variants, annotation matrix *A* ∈ *R*^*S*×*L*^ encoding *L* functional annotations, and covariate matrix *F* ∈ *R* ^*N*×*K*^ for *K* factors (e.g., age, sex and leading genetic principal components), gene-level burden tests consider the following linear model (**Figure 1a**):

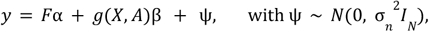

where *g*(*X, A*) = [*g*(*x, A*), …, *g*(*x*_1_, *A*)] ∈ *R* ^*N*×1^ is a function aggregating variant effects for each individual based on annotations *A* in a gene-level burden score, *α* ∈ *R*^*K*^ is the vector of covariate effects, β is the effect size of the burden score, and 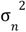 is the residual variance. For example, *g*(*X, A*) could be the sum of putative loss-of-function (pLoF) variants within a gene for each individual (Cirulli et al. 2020). Within this framework, statistical association between the gene burden score and the phenotype is assessed by testing whether β≠0.

**Figure 1.**
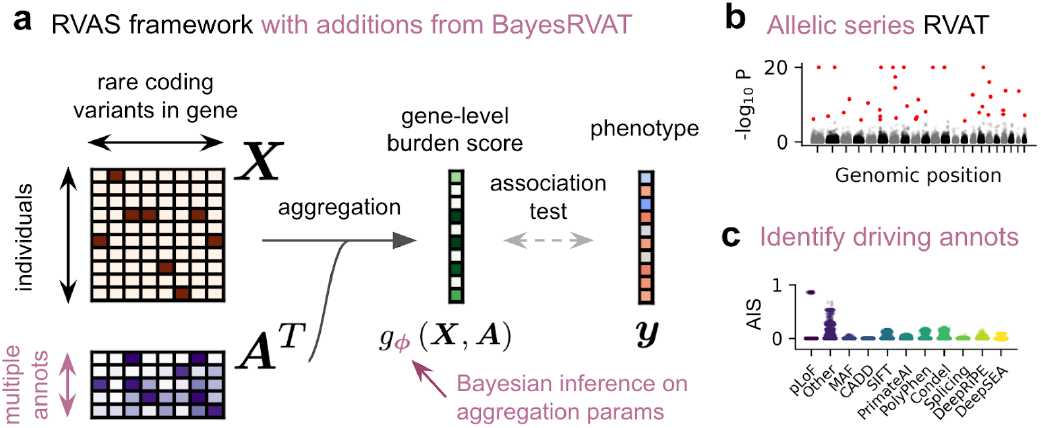
Overview of the BayesRVAT framework. (**a**) In rare variant association tests (RVAT), rare variants *X* and their annotations *A* are aggregated into a gene burden score, which is tested for association with the phenotype *y*. BayesRVAT explicitly introduces aggregation function *g*_ϕ_ (*X, A*) and a prior over aggregation parameters ϕ. (**b**) BayesRVAT enables scalable gene burden testing accounting for multiple annotations. (**c**) It also provides annotation importance scores (AIS) for each analyzed gene-trait pair.

In BayesRVAT, we enhance flexibility by making the aggregation function *g*_ϕ_ (*X, A*) dependent on parameters ϕ, which describe how variants are integrated into a burden score based on their annotations (**Figure 1a**). Specifically, we use a linear model with saturation:

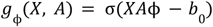

where annotations *A* are processed such that higher values correspond to more deleterious effects (**Methods**), *b*_0_ shifts the input such that individuals without rare variants have scores close to zero, and the sigmoid function σ accounts for saturation effects, where additional variants do not further increase the burden once the gene function is already lost. We introduce priors on ϕ to reflect biological expectations while modeling uncertainty: we introduce strong effect priors for pLoF variants, while considering weaker effect and higher variance priors for other annotations (**Methods, Supplementary figure A1**).

We employ variational inference to estimate posterior distributions over ϕ and estimate model parameters for each gene-trait pair (Ranganath et al. 2014; Rezende et al. 2014; Kingma 2013) (**Methods**). After the estimation step, BayesRVAT provides a gene-level association P value (**Methods**) and Annotation Importance Scores (AIS), which quantify the extent to which specific annotations drive the association for the analyzed gene-trait pair (**Methods**).

### Simulations

We evaluated BayesRVAT using simulated data from unrelated individuals in the UK Biobank (UKB) cohort (**Methods**). Briefly, we simulated gene-level genetic effects from real variant data using a saturated additive model, and varied key parameters, including sample size, variance explained by the burden score, and the number of contributing annotations (**Methods**). For variant annotations, we considered 25 features, including three derived from variant consequences, allele frequency, five functional impact scores, two splicing prediction scores, eight RNA-binding propensities, and six regulatory annotations (**Methods**).

First, we assessed the statistical calibration of BayesRVAT when simulating under a null model with no genetic effects, which yielded calibrated P values across different simulated sample sizes (**Figure 2a, Supplementary Figure A2**). Next, we compared the statistical power of BayesRVAT against commonly used burden tests, including classical pLoF burden testing; ACAT-Conseq, which aggregates separate tests for pLoF, missense, and other non-synonymous variants (ACAT-Conseq, **Methods**), corresponding to the basic allelic series burden test in (McCaw et al. 2023); and ACAT-MultiAnnot, which aggregates separate tests for each of the 25 analyzed annotations (**Methods**), similar to the method in (Li et al. 2020). Notably, BayesRVAT consistently outperformed alternative methods, sustaining higher power in more complex genetic architectures with increasing contributing annotations while remaining robust to non-informative annotations (**Figure 2b**). The superior performance of BayesRVAT was maintained across varying sample sizes and levels of variance explained by the burden score (**Supplementary Figure A3**).

**Figure 2.**
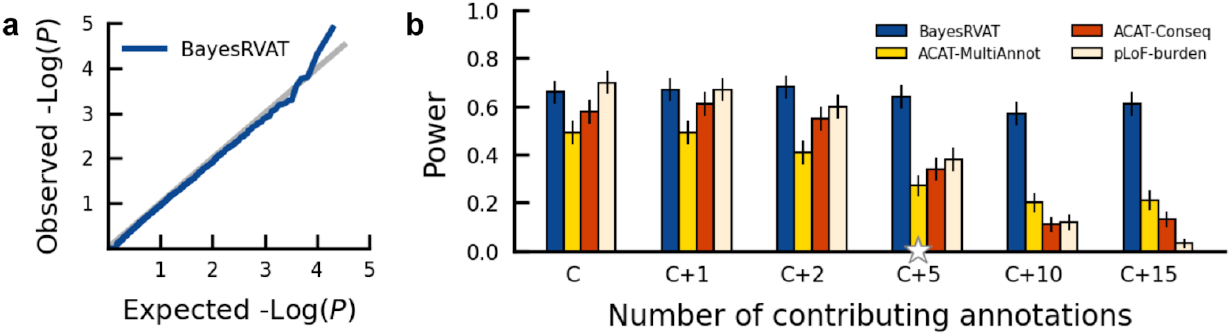
Evaluation of calibration and power in BayesRVAT using synthetic data. (**a**) QQ plot assessing the calibration of P values from BayesRVAT on synthetic data generated under the null model with no genetic effects. (**b**) Statistical power comparison between BayesRVAT, ACAT-MultiAnnot, ACAT-Conseq, and pLoF-burden test, across varying numbers of contributing continuous annotations: simulating only effects from pLoF and missense consequences (C), and considering additional effects from 1 (C+1), 2 (C+2), 5 (C+5), 10 (C+10), and 15 continuous annotations (C+15; **Methods**). Power is measured at the exome-wide significance threshold of *P* < 2. 5 × 10 ^−6^, computed over 100 replicates for each scenario.

We further benchmarked BayesRVAT against state-of-the-art variance component tests and optimal methods integrating burden and variance component testing, such as SKAT-O (Lee et al. 2012). Notably, BayesRVAT maintained superior power over both approaches (**Supplementary Figure A4**).

### Analysis of blood biomarkers

We applied BayesRVAT and alternative burden test strategies to analyze twelve blood traits from the UKB cohort, considering the same set of 25 annotations used in simulations (**Methods**). BayesRVAT identified a greater number of significant gene-trait associations compared to other methods (130 for BayesRVAT vs 118 for ACAT-MultiAnnot, 92 for ACAT-Conseq, and 86 for pLoF; Bonferroni-adjusted *P* < 5 × 10^−2^ ; **Figure 3a-b, Supplementary Figure A5**), also showing well-calibrated P values under genotype permutation tests (**Figure 3c**). Interestingly, BayesRVAT consistently outperformed ACAT-MultiAnnot (**Figure 3b, Supplementary Figure A5**), except when allelic series assumptions were violated. This occurs when other annotations have stronger effects than pLoF, directly contradicting the priors in BayesRVAT (**Supplementary Figure A6**). Notably, compared to the widely used SKAT variance component and SKAT-O optimal tests (Lee et al. 2012; Wu et al. 2011), BayesRVAT also identified more associations (**Supplementary Figure A7**).

**Figure 3.**
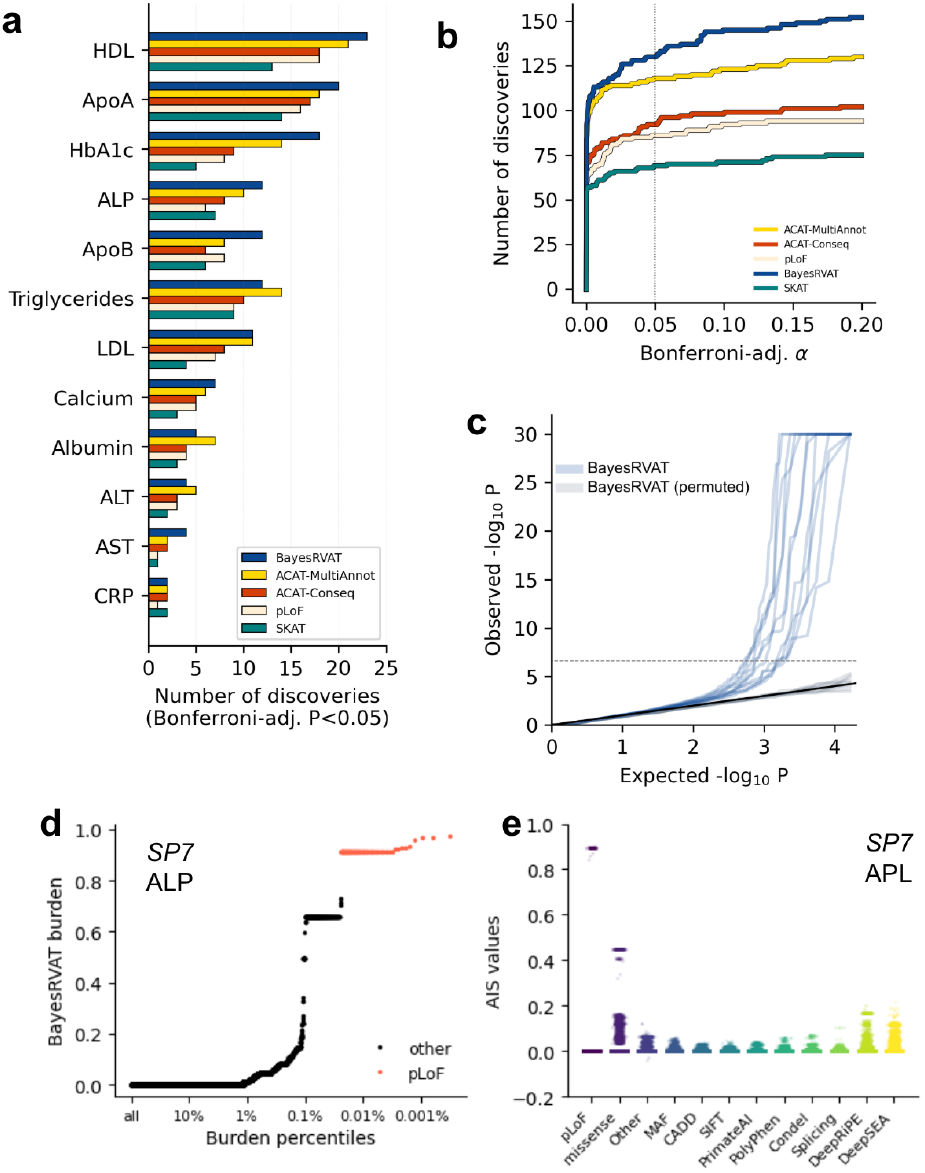
Analysis of blood biomarkers in the UK Biobank. (**a**) Number of significant gene-trait associations (Bonferroni-adjusted *P* < 5 × 10^−2^) discovered by BayesRVAT, ACAT-MultiAnnot, ACAT-Conseq, and pLoF burden tests for each analyzed blood trait. (**b**) Cumulative number of discoveries at varying Bonferroni-adjusted significance thresholds α. (**c**) QQ plot showing the distribution of P values from BayesRVAT in real data and under a null with permuted genotype data, confirming well-calibrated P values. (**d**) Burden scores learned by BayesRVAT for *SP7* and ALP across burden percentiles, showing individuals carrying pLoF mutations in red. (**e**) Annotation importance scores (AIS) from BayesRVAT for the association between *SP7* and ALP, which highlights contributions from missense, DeepRiPE, and DeepSEA annotations.

Among the associations uniquely identified by BayesRVAT, several showed strong biological relevance (**Figure 3d-e, Supplementary Figure A8**). For instance, BayesRVAT uniquely detected an association between *SP7* and alkaline phosphatase (ALP). *SP7* is a transcription factor, which was shown before to regulate the expression of ALP (Lui et al. 2022; Yoshida et al. 2012). In this case, BayesRVAT’s burden score assigned higher weight to annotations beyond loss-of-function (pLoF) mutations (**Figure 3d**), with AIS scores indicating contributions from missense, DeepRiPE, and DeepSEA annotations

### Optimal test integrating BayesRVAT burden with variance component tests

Next, we assessed the performance of BayesRVAT when combined with variance component models and compared it to other optimal tests integrating both burden and variance component models, such as SKAT-O (Lee et al. 2012). Specifically, we compared BayesRVAT + SKAT against pLoF + SKAT, ACAT-Conseq + SKAT, and ACAT-MultiAnnot + SKAT (**Methods**). Notably, even without SKAT integration, BayesRVAT outperformed all optimal tests (130 for BayesRVAT vs 120 for ACAT-MultiAnnot + SKAT, 99 for ACAT-Conseq + SKAT, and 101 for pLoF + SKAT; Bonferroni-adjusted α = 0. 05; **Figure 4, Supplementary Figure A7**). Adding SKAT to BayesRVAT provided a modest improvement, identifying three additional significant associations (Bonferroni-adjusted α = 0. 05; **Figure 4**).

**Figure 4.**
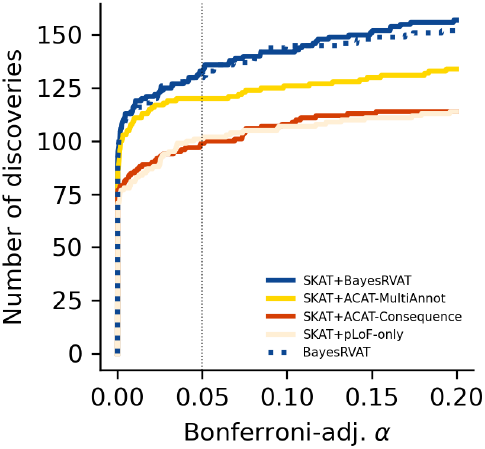
Integration of BayesRVAT and other burden tests with variance component models. Comparison of cumulative significant gene-trait discoveries (Bonferroni-adjusted *P* < 5 × 10^−2^) across various methods: BayesRVAT + SKAT, ACAT-MultiAnnot + SKAT, ACAT-Conseq + SKAT, and pLoF + SKAT. We also report BayesRVAT without SKAT integration, highlighting its superior performance compared to other optimal tests even without SKAT integration.

### Application to disease traits

We applied BayesRVAT to analyze eight disease traits with binary notations: type 2 diabetes, atrial fibrillation, coronary artery disease, asthma, hypertension, age-related macular degeneration (AMD) and other retinal diseases, glaucoma, and cataract (**Methods**). For comparison, we also evaluated logistic regression-based burden tests using pLoF, ACAT-Conseq, and ACAT-MultiAnnot.

BayesRVAT consistently identified more associations than alternative burden tests, detecting ten significant gene-trait pairs (Bonferroni-adjusted P < 0.05) compared to seven for ACAT-MultiAnnot, four for ACAT-Conseq, and seven for pLoF (Bonferroni-adjusted α < 5 × 10^−2^; **Figure 5a**; **Supplementary Table A1**), underscoring its sensitivity. Furthermore, at these loci, BayesRVAT assigned stronger statistical evidence to associations, as reflected in the distribution of P values (**Figure 5b**).

**Figure 5.**
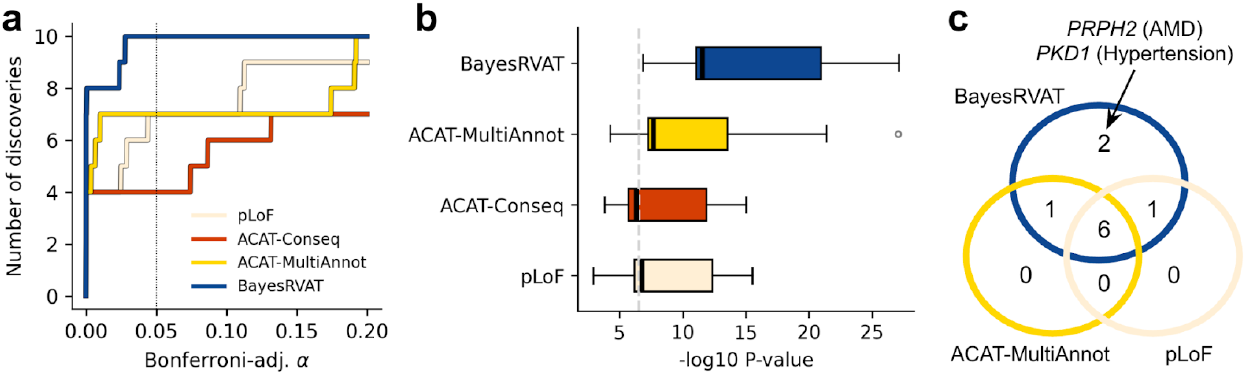
BayesRVAT outperforms alternative burden tests in disease trait analyses. (**a**) Step function showing the cumulative number of significant gene-trait associations as a function of Bonferroni-adjusted significance threshold α. BayesRVAT (blue) consistently identifies more associations than pLoF (beige), ACAT-Conseq (red), and ACAT-MultiAnnot (yellow) across all thresholds. (**b**) Distribution of P values for the nine significant associations identified across all methods (Bonferroni-adjusted P < 0.05 for at least one method). (**c**) Venn diagram illustrating the overlap of significant gene-trait associations detected by each method. BayesRVAT recovers all signals found by other methods and uniquely identifies associations between *PKD1* and hypertension, and between *PRPH2* and AMD and other retinal diseases.

BayesRVAT uniquely identified two associations not detected by any other method. The first is an association between *PKD1* and hypertension (**Figure 5c**), which can be detected at larger sample sizes (Karczewski et al. 2022). The second is a link between *PRPH2* and AMD and other retinal diseases (**Figure 5c**). *PRPH2* encodes a structural protein in the outer segments of retinal photoreceptors (Kalaw et al. 2025) and multiple *PRPH2* variants have been implicated in retinal conditions (AlAshwal et al. 2025).

## Discussion

In this work, we introduce BayesRVAT, a flexible Bayesian framework for rare variant association testing that integrates multiple functional annotations to improve gene-trait association discovery. By modeling how different annotations contribute to gene disruption and ultimately affect phenotype, BayesRVAT models allelic series, enabling a data-driven aggregation of effects for each analyzed gene-phenotype pair. This is important because different annotations may contribute to gene function disruption in ways that vary across genes and phenotypes (Ritchie and Flicek 2015).

Importantly, the Bayesian allelic series framework in BayesRVAT is particularly relevant given the rapid expansion of machine learning models trained on biological sequences to predict protein structure (Cheng et al. 2023; Gao et al. 2023), gene expression (Avsec et al. 2021; Linder et al. 2023), splicing (Jaganathan et al. 2019; Wagner et al. 2023), as well as through self-supervised learning approaches (Benegas et al. 2024; Dalla-Torre et al. 2023; Rives et al. 2021; Nguyen et al. 2024). These models have generated large-scale variant pathogenicity estimates, necessitating new methods that integrate these predictions within allelic series models, a gap addressed by BayesRVAT.

BayesRVAT consistently outperformed conventional burden tests, achieving higher power when genetic effects align with its allelic series assumptions. In real data applications, BayesRVAT detected 10.2% more significant associations in blood biomarker analyses and uncovered additional gene-trait associations in disease studies missed by other methods. For example, BayesRVAT uniquely identified the association between *SP7* and alkaline phosphatase (ALP), consistent with the SP7’s role in ALP expression (Lui et al. 2022; Yoshida et al. 2012). It also detected associations between *EPB42* and glycated hemoglobin (**Supplementary Figure A8**), in line with recent findings linking *EPB42* variants to glycemic traits (Kim et al. 2022), and between *NPC1L1* and apolipoprotein B (**Supplementary Figure A8**), reflecting its impact on lipid transport and metabolism (Jia et al. 2011). In the disease trait GWAS, BayesRVAT uniquely linked *PRPH2* to AMD and other retinal diseases. Previously associated with retinitis pigmentosa and pattern dystrophies (AlAshwal et al. 2025), *PRPH2* may contribute to AMD susceptibility through mechanisms affecting photoreceptor stability and function.

We further note that BayesRVAT has limitations that future work can address. First, although BayesRVAT’s runtime scales linearly with cohort size (**Supplementary Figure A9**), making it feasible for large biobank datasets, it remains more computationally demanding than standard burden tests due to the need for variational inference. To mitigate this, we propose a two-stage strategy where BayesRVAT is run selectively on genes showing preliminary evidence of association in simpler burden tests, significantly reducing computational burden (**Methods**).

Second, BayesRVAT does not explicitly model interactions between variant annotations due to the simple form of its aggregation function. Additionally, while our choice of priors extends the pLoF test, allowing other annotations to contribute to partial gene disruption, alternative formulations may be explored. Future work may incorporate interaction features between annotations or embeddings from self-supervised DNA and protein models to capture richer functional and structural dependencies (Benegas et al. 2024; Dalla-Torre et al. 2023; Rives et al. 2021; Nguyen et al. 2024).

Finally, BayesRVAT is currently limited to analysis in cohorts of unrelated individuals. Looking forward, we aim to expand BayesRVAT to incorporate mixed-model adjustments with sparse kinship matrices, which can improve rare variant testing in structured and related cohorts (Zhou et al. 2022). More broadly, ongoing efforts aim to facilitate integration with cloud-based genomic analysis platforms, such as the UK Biobank Research Analysis Platform, to enhance accessibility and scalability for large-scale rare variant studies.

## Methods

### A Bayesian Framework for RVAT

#### The burden test framework

Gene-level burden testing is performed using the following linear model

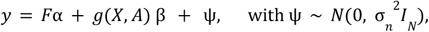

where *y* is the phenotype vector (*N* × 1) for *N* individuals, *F* the covariate matrix (*N* × *K*) for *K* covariates, and α the vector of covariate effects (*K* × 1). The function *g*(*X, A*) computes a gene-level burden score (*N* × 1), by aggregating the variant matrix *X* (*N* × *S*), where *S* is the number of rare variants, using the annotation matrix *A* (*S* × *L*), where *L* is the number of annotations per variant. The coefficient β represents the effect of the burden test, while *ψ* is the residual error, assumed to follow a normal distribution with variance 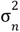. A classical choice for *g*(*X, A*) is the sum of pLoF variants within a gene for each individual (Cirulli et al. 2020). Within this framework, statistical association between the gene burden score and the phenotype is assessed by testing whether β≠0.

#### Bayesian formulation

In BayesRVAT, we parameterize the aggregation function *g*_ϕ_ (*X, A*) with parameters ϕ and introduce a prior distribution *p*(ϕ) that incorporates our prior beliefs on how to aggregate variants into a burden score based on their annotations (**Figure 1a**).

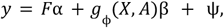

Introducing compact notations for input data *D* = {*F, X, A*} and model parameters θ = {α, β, σ^2^}, the model marginal likelihood can be written as

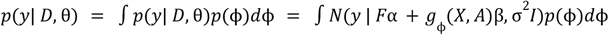

We note that our Bayesian framework allows the data for each gene and trait to update these prior beliefs, effectively adapting the posterior on aggregation parameters *p*(ϕ) to the specific gene/trait pair being analyzed.

#### Choice of aggregation function and priors

After preprocessing annotations *A* to ensure that higher values correspond to more deleterious effects, we assume linear variant effects in *A* and use an additive model with saturation to collapse the contributions of multiple variants into a single gene burden score, *g*_ϕ_ (*X, A*) = *σ*(*XA*ϕ − *b*_0_). Here, *b*_0_ is a bias term that ensures individuals carrying no rare variants receive a burden score close to zero, and the sigmoid function introduces a saturation mechanism—once the gene is impaired, additional variants no longer contribute to disruption. Our priors on ϕ reflect biological knowledge (**Supplementary Figure A1**), setting pLoF effects’ prior such that carriers are highly likely to receive a gene burden score close to one. In contrast, for other non-synonymous variants, we use weaker priors with greater variability, accounting for the uncertainty on their effects. For functional, regulatory and splicing annotation scores, we apply priors that allow moderate, positive adjustments to the burden score (**Supplementary Figure A1**).

#### Continuous and binary phenotypes

BayesRVAT supports both continuous and binary traits by adapting the phenotype likelihood function accordingly. For continuous traits, we assume a normal likelihood (as described above). For case-control traits, we use a Bernoulli likelihood with a sigmoid link function to map the linear predictor (including covariate and gene burden effects) to the probability of case status. While all model derivations are presented in the continuous trait case, the adaptation to binary traits follows directly by substituting the likelihood function.

#### Optimization

The optimization of model parameters θ by maximum likelihood is intractable for a general aggregation function *g*_ϕ_ (*X, A*) due to the integral over ϕ. We thus use black-box variational inference (Ranganath et al. 2014), which approximates the true posterior *p*(ϕ |*y, D*, θ) with a simpler variational distribution *q*_*ψ*_ (ϕ) parameterized by *ψ*. Within this framework, we optimize both the model parameters θ and the variational parameters ψ by maximizing the Evidence Lower Bound (ELBO):

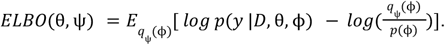

We assume a mean-field Gaussian variational posterior for *q*_*ψ*_ (ϕ) and approximate the expectation using Monte Carlo sampling (Rezende et al. 2014; Kingma 2013). By maximizing the ELBO, we jointly estimate the values of 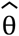 and the variational parameters 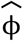, providing approximations of the maximum likelihood estimator of θ and the exact posterior distribution, respectively.

#### Association testing

Within the BayesRVAT framework, we can assess associations between gene burden scores and trait values by testing the hypothesis β ≠ 0 (**Figure 1b**). As the likelihood ratio test statistic is intractable due to the integral over ϕ, in the alternative hypothesis, we introduce an approximate Likelihood Ratio Test statistic. Briefly, we replace the intractable log marginal likelihood under the alternative hypothesis with the importance-weighted variational evidence lower bound (IW-ELBO) (Burda et al. 2015):

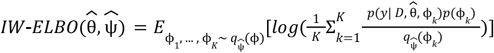

which is computed using the approximate maximum likelihood estimators 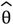 and the variational posterior 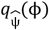, obtained through optimizing the ELBO. The IW-ELBO is a tighter bound on the log marginal likelihood compared to the standard ELBO, leading to more accurate P values. We note that this approximation yields conservative P values as it replaces the log marginal likelihood under the alternative hypothesis with its lower bound, while the log marginal likelihood under the null hypothesis can be computed exactly. As a result, test statistics are lower than in an exact likelihood ratio test. Increasing the number of importance samples (K) improves the accuracy of P values (see **Supplementary Figure A10**).

#### Annotation Importance Scores

Similar to sensitivity analysis (Iooss and Lemaître 2015), we can evaluate the importance of a set of annotations by comparing the gene burden scores computed using all annotations (denoted as *A*_1_) with the scores obtained by setting a subset of annotations to their median values (denoted as *A*_0_). Specifically, we compute the expected value of the difference between these two burden scores:

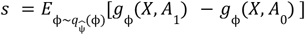

The result *s* ∈ *R* ^*N*×1^ quantifies how much each individual’s gene burden is influenced by the annotations under investigation. We refer to these scores as Annotation Importance Scores (AIS, **Figure 1c**).

#### Implementation Details

We implemented BayesRVAT and its derivatives using Numpy and SciPy, optimizing the ELBO using the L-BFGS in *scipy*.*optimize* (Byrd et al. 1995; Zhu et al. 1997; Morales and Nocedal 2011). Since L-BFGS requires noise-free loss and gradient evaluations, we fixed the Monte Carlo samples to approximate the expectation in the ELBO during its optimization, following (Loh et al. 2015), and used 16 Monte Carlo samples. For association testing, P values were computed by approximating the IW-ELBO using 30 Monte Carlo estimates, each based on 16 importance samples.

### Preprocessing of the UK BioBank dataset

All experiments were conducted using the UKB cohort (Bycroft et al. 2018) based on the latest whole-exome sequencing (WES) release. Individual and variant quality control (QC) followed the protocols from GeneBass (Karczewski et al. 2022). Variant annotation was performed using the pipeline described in (Clarke et al. 2024). The final processed dataset includes 329,087 unrelated European individuals, 5,845,828 variants with MAF < 0.1%, 16,458 genes, and 25 variant annotations.

#### Genetic data QC

We closely followed (Karczewski et al. 2022) for variant QC. For sample QC, we followed the procedure used in (Neale 2018). Briefly, we applied filters to remove samples flagged as used in PCA calculations (i.e., unrelated samples) and those with sex chromosome aneuploidy. To restrict the dataset to individuals of British ancestry, we utilized the provided principal components (PCs), selecting individuals within 7 standard deviations from the first six PCs, and filtered to self-reported ethnicities, specifically ‘white-British,’ ‘Irish,’ or ‘White’. To account for batch effects in whole-exome sequencing (WES), we followed a similar approach to (Karczewski et al. 2022). Briefly, we assessed the coverage around genes and used Scanpy (Wolf et al. 2018) to cluster individuals based on 8 PCs of the coverage (10 nearest neighbours, Leiden clustering with resolution of 1). Smaller outlying clusters were excluded, and WES batch clusters were inferred using the Leiden clustering at a lower resolution (resolution 0.1), identifying three main groups. These cluster labels were used as covariates in downstream association studies to mitigate any potential batch-related bias (Li et al. 2023; Law et al. 2014).

#### Variant Annotation

We defined consequences using VEP (McLaren et al. 2016), and classified pLoF as any splice donor, frameshift, splice acceptor, stop-gained, stop-lost, or start-lost variant. Our total set of 25 annotations consists of three consequence-based annotations consequences (pLoF, missense, and others), minor allele frequency, CADD (Rentzsch et al. 2019), SIFT (Kumar et al. 2009), PolyPhen-2 (Adzhubei et al. 2010), PrimateAI (Sundaram et al. 2018), Condel (González-Pérez and López-Bigas 2011), SpliceAI delta score (Jaganathan et al. 2019) and the AbSpliceDNA score (Wagner et al. 2023), eight RNA-binding protein binding propensity delta scores derived from DeepRiPE (Ghanbari and Ohler 2020), and six regulatory scores, derived as principal components of DeepSEA delta embeddings (Zhou and Troyanskaya 2015). We ensured that for any annotation, higher values correspond to higher pathogenicity. For example, we used 1 − *SIFT* rather than *SIFT*, and built the MAF-based annotation as 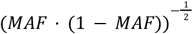. Finally, we used the PHRED scale to ensure consistency across annotations, i.e., − 10*log*_10_ (*rank*(− *score*)/*L*) where *L* is the total number of variants (Li et al. 2020).

### Benchmarking Against Alternative Burden Testing Methods

To assess the performance of BayesRVAT compared to other commonly used burden testing techniques, we benchmarked our method against: pLoF-Burden, a burden test based on the sum of pLoF variants; ACAT-Conseq, which performs separate association tests on the sums of pLoF, missense, and other non-synonymous variants, with P values aggregated using the Aggregated Cauchy Association Test (Liu et al. 2019)—a strategy similar to the simple allelic series models considered in (McCaw et al. 2022); and ACAT-MultiAnnot, which runs burden tests across each of the annotations used in BayesRVAT, combining results using ACAT, similar to the approach in (Li et al. 2020). All burden tests were implemented as likelihood ratio tests within a linear model framework, adjusting for age, sex, the top twenty genetic principal components, and WES batch effect covariates. For continuous traits (Gaussian likelihood), maximum likelihood estimates have a closed-form solution, whereas for binary traits (Bernoulli likelihood), we optimize the null and alternative models using L-BFGS, implemented in SciPy.

### Simulations

We used synthetic data to assess both the calibration and power of BayesRVAT under various simulated conditions, based on 100,000 individuals from the processed UKB cohort. To evaluate power, we simulated additive genetic effects from a gene burden test using the additive model with saturation. We varied key parameters such as sample size, variance explained by the burden, and the number of continuous annotations contributing to the burden score. Power was estimated at the exome-wide significance threshold of *P* < 2. 5 × 10^−6^, with 100 replicates performed for each simulation configuration. To assess calibration, we simulated phenotypes under a null model with no genetic effects. Full details on the simulation procedures are provided in **Supplementary Information**.

### Real data analyses in UK Biobank

#### Blood biomarkers

We applied BayesRVAT and baselines to exome sequencing data from the UK Biobank, selecting unrelated individuals of self-reported White British ancestry. We analyzed twelve key blood biomarkers: LDL cholesterol, HDL cholesterol, apolipoprotein A, apolipoprotein B, triglycerides, glycated hemoglobin, alanine aminotransferase, aspartate aminotransferase, albumin, alkaline phosphatase, calcium, and C-reactive protein. To ensure normality, all biomarker phenotypes were transformed using a rank-based inverse normal transformation.

#### Integration with SKAT

We used the classical SKAT variance component test and combined its results with each burden test using the Aggregated Cauchy Association Test (ACAT), forming an optimal test. Note that in comparison with SKAT-O (**Supplementary Figures A4, A7**), we used the standard SKAT-O implementation in R (Lee et al. 2012) rather than this ACAT-based equivalent.

#### Disease code analyses

We analyzed eight disease traits: type 2 diabetes (*fieldID*=130708; 26,328 cases, 302,759 controls), asthma (*fieldID*=131494; 47,125 cases, 281,962 controls), coronary artery disease (*fieldID*=131306; 33,878 cases, 295,209 controls), age-related macular degeneration (*fieldID*=131182; 11,083 cases, 318,004 controls), glaucoma (*fieldID*=131186; 12,633 cases, 316,454 controls), cataract (*fieldID*=131166; 30,566 cases, 298,521 controls), and atrial fibrillation (*fieldID*=131350; 25,813 cases, 303,274 controls). These conditions are relatively common (≥5,000 cases in UK Biobank) and span respiratory, metabolic, cardiovascular, and ocular diseases. All traits were derived from “Date first reported” phenotypes, curated by UK Biobank (UKB) using ICD codes, primary care records, and self-reported data. For each trait, we tested 16,017 genes, yielding a total of 128,136 gene-trait association tests. Statistical significance was defined using a Bonferroni-corrected P < 0.05 (raw P < 3.9×10^−7^).

### Computational complexity and run time

BayesRVAT exhibits linear computational complexity with respect to sample size, making it feasible for large biobank datasets. To empirically assess runtime scaling, we measured the average time to fit a single gene at different cohort sizes in simulations (**Supplementary Figure A10**), yielding 11 ± 0. 42 seconds for N=50,000, 29. 32 ± 1. 13 seconds for N=100,000, 71. 1 ± 2. 44 seconds for N=200,000, and 112. 1 ± 3. 17 seconds for N = 300,000 individuals. While BayesRVAT remains computationally tractable, its runtime is substantially higher than standard burden tests—e.g., ACAT-MultiAnnot requires 0. 12 ± 0. 01 for N = 300,000. To reduce computational burden, we propose a two-stage filtering strategy, applying BayesRVAT selectively to genes showing preliminary association signals in simpler burden tests. Since most genes are not associated with the phenotype and yield approximately uniform P-values under the null, applying a pre-filtering threshold of P<10^−2^ reduces the number of genes analyzed by 90%, cutting total runtime to 5 hours for N = 300,000. A stricter threshold of P<10^−3^ would further reduce the runtime to just a few hours for N = 300,000. All run times were estimated on Intel Xeon Gold 6134 CPUs with 32 logical cores.

### Use of Artificial Intelligence

In the preparation of this manuscript, we utilized the large language model GPT-4 (https://chat.openai.com/) for editing assistance, including language polishing and clarification of text. While this tool assisted in refining the manuscript’s language it was not used to generate contributions to the original research, data analysis, or interpretation of results. All final content decisions and responsibilities rest with the authors.

## Acknowledgements

The authors would like to thank Brian Clarke, Julien Gagneur, Eva Holtkamp, Shubhankar Londhe and Oliver Stegle for helpful discussions. This research has been conducted using the UK Biobank Resource (Application Number 87065). F.P.C. and A.N. were funded by the Free State of Bavaria’s Hightech Agenda through the Institute of AI for Health (AIH). L.S. acknowledges the support of Friedrich-Alexander-Universität Erlangen-Nürnberg. A.N. and L.S. acknowledge the Helmholtz Association under the joint research school “Munich School for Data Science - MUDS”.

## Authors’ contributions

A.N. and F.P.C. implemented the methods. A.N., L.S., and F.P.C. analyzed the data. A.N. and F.P.C. wrote the manuscript, with all authors contributing to revisions. T.K. provided critical input on methodological development, including variational inference and optimization. All authors contributed to the interpretation of the results. F.P.C. conceived the study with support from N.C. F.P.C. and N.C. supervised the work.

## References

Adzhubei IA, Schmidt S, Peshkin L, Ramensky VE, Gerasimova A, Bork P, Kondrashov AS, Sunyaev SR. 2010. A method and server for predicting damaging missense mutations. Nature methods 7: 248–249.

AlAshwal SM, Yassin SH, Kalaw FGP, Borooah S. 2025. Prph2-associated retinal diseases: A systematic review of phenotypic findings. Am J Ophthalmol 271: 7–30.

Avsec Ž, Agarwal V, Visentin D, Ledsam JR, Grabska-Barwinska A, Taylor KR, Assael Y, Jumper J, Kohli P, Kelley DR. 2021. Effective gene expression prediction from sequence by integrating long-range interactions. Nat Methods 18: 1196–1203.

Backman JD, Li AH, Marcketta A, Sun D, Mbatchou J, Kessler MD, Benner C, Liu D, Locke AE, Balasubramanian S, et al. 2021. Exome sequencing and analysis of 454,787 UK Biobank participants. Nature 599: 628–634.

Benegas G, Albors C, Aw AJ, Ye C, Song YS. 2024. GPN-MSA: an alignment-based DNA language model for genome-wide variant effect prediction. bioRxivorg. http://dx.doi.org/10.1101/2023.10.10.561776.

Brandes N, Goldman G, Wang CH, Ye CJ, Ntranos V. 2023. Genome-wide prediction of disease variant effects with a deep protein language model. Nat Genet 55: 1512–1522.

Brandes N, Linial N, Linial M. 2020. PWAS: proteome-wide association study—linking genes and phenotypes by functional variation in proteins. Genome biology 21: 173.

Burda Y, Grosse R, Salakhutdinov R. 2015. Importance weighted autoencoders. arXiv preprint arXiv:1509 00519.

Byrd RH, Lu P, Nocedal J, Zhu C. 1995. A limited memory algorithm for bound constrained optimization. SIAM J Sci Comput 16: 1190–1208.

Cheng J, Novati G, Pan J, Bycroft C, Žemgulytė A, Applebaum T, Pritzel A, Wong LH, Zielinski M, Sargeant T, et al. 2023. Accurate proteome-wide missense variant effect prediction with AlphaMissense. Science 381: eadg7492.

Cirulli ET, White S, Read RW, Elhanan G, Metcalf WJ, Tanudjaja F, Fath DM, Sandoval E, Isaksson M, Schlauch KA, et al. 2020. Genome-wide rare variant analysis for thousands of phenotypes in over 70,000 exomes from two cohorts. Nat Commun 11: 542.

Clarke B, Holtkamp E, Öztürk H, Mück M, Wahlberg M, Meyer K, Munzlinger F, Brechtmann F, Hölzlwimmer FR, Lindner J, et al. 2024. Integration of variant annotations using deep set networks boosts rare variant association testing. Nat Genet 56: 2271–2280.

Dalla-Torre H, Gonzalez L, Revilla JM, Carranza NL, Grzywaczewski AH, Oteri F, Dallago C, Trop E, Sirelkhatim H, Richard G, et al. 2023. The Nucleotide Transformer: Building and evaluating robust foundation models for human genomics. http://dx.doi.org/10.1101/2023.01.11.523679.

Gao H, Hamp T, Ede J, Schraiber JG, McRae J, Singer-Berk M, Yang Y, Dietrich ASD, Fiziev PP, Kuderna LFK, et al. 2023. The landscape of tolerated genetic variation in humans and primates. Science 380: eabn8153.

Ghanbari M, Ohler U. 2020. Deep neural networks for interpreting RNA-binding protein target preferences. Genome research 30: 214–226.

González-Pérez A, López-Bigas N. 2011. Improving the assessment of the outcome of nonsynonymous SNVs with a consensus deleteriousness score, Condel. The American Journal of Human Genetics 88: 440–449.

Iooss B, Lemaître P. 2015. A review on global sensitivity analysis methods. Uncertainty management in simulation-optimization of complex systems: algorithms and applications 101–122.

Jaganathan K, Panagiotopoulou SK, McRae JF, Darbandi SF, Knowles D, Li YI, Kosmicki JA, Arbelaez J, Cui W, Schwartz GB, et al. 2019. Predicting splicing from primary sequence with deep learning. Cell 176: 535–548.

Jia L, Betters JL, Yu L. 2011. Niemann-pick C1-like 1 (NPC1L1) protein in intestinal and hepatic cholesterol transport. Annual review of physiology 73: 239–259.

Jurgens SJ, Choi SH, Morrill VN, Chaffin M, Pirruccello JP, Halford JL, Weng L-C, Nauffal V, Roselli C, Hall AW, et al. 2022. Analysis of rare genetic variation underlying cardiometabolic diseases and traits among 200,000 individuals in the UK Biobank. Nature genetics 54: 240–250.

Kalaw FGP, Wagner NE, de Oliveira TB, Everett LA, Yang P, Pennesi ME, Borooah S. 2025. Using multimodal imaging to refine the phenotype of PRPH2-associated retinal degeneration. Ophthalmol Retina 9: 69–77.

Karczewski KJ, Solomonson M, Chao KR, Goodrich JK, Tiao G, Lu W, Riley-Gillis BM, Tsai EA, Kim HI, Zheng X, et al. 2022. Systematic single-variant and gene-based association testing of thousands of phenotypes in 394,841 UK Biobank exomes. Cell Genomics 2.

Kim YJ, Moon S, Hwang MY, Han S, Jang H-M, Kong J, Shin DM, Yoon K, Kim SM, Lee J-E, et al. 2022. The contribution of common and rare genetic variants to variation in metabolic traits in 288,137 East Asians. Nature communications 13: 6642.

Kingma DP. 2013. Auto-encoding variational bayes. arXiv preprint 1312-6114.

Kumar P, Henikoff S, Ng PC. 2009. Predicting the effects of coding non-synonymous variants on protein function using the SIFT algorithm. Nature protocols 4: 1073–1081.

Law CW, Chen Y, Shi W, Smyth GK. 2014. voom: Precision weights unlock linear model analysis tools for RNA-seq read counts. Genome biology 15: 1–17.

Lee S, Wu MC, Lin X. 2012. Optimal tests for rare variant effects in sequencing association studies. Biostatistics 13: 762–775.

Li B, Leal SM. 2008. Methods for detecting associations with rare variants for common diseases: application to analysis of sequence data. The American Journal of Human Genetics 83: 311–321.

Linder J, Srivastava D, Yuan H, Agarwal V, Kelley DR. 2023. Predicting RNA-seq coverage from DNA sequence as a unifying model of gene regulation. Genomics. https://www.biorxiv.org/content/10.1101/2023.08.30.555582v1.

Li T, Zhang Y, Patil P, Johnson WE. 2023. Overcoming the impacts of two-step batch effect correction on gene expression estimation and inference. Biostatistics 24: 635–652.

Liu Y, Chen S, Li Z, Morrison AC, Boerwinkle E, Lin X. 2019. ACAT: a fast and powerful p value combination method for rare-variant analysis in sequencing studies. The American Journal of Human Genetics 104: 410–421.

Li X, Li Z, Zhou H, Gaynor SM, Liu Y, Chen H, Sun R, Dey R, Arnett DK, Aslibekyan S, et al. 2020. Dynamic incorporation of multiple in silico functional annotations empowers rare variant association analysis of large whole-genome sequencing studies at scale. Nature genetics 52: 969–983.

Loh P-R, Bhatia G, Gusev A, Finucane HK, Bulik-Sullivan BK, Pollack SJ, Schizophrenia Working Group of Psychiatric Genomics Consortium, de Candia TR, Lee SH, Wray NR, et al. 2015. Contrasting genetic architectures of schizophrenia and other complex diseases using fast variance-components analysis. Nat Genet 47: 1385–1392.

Lui JC, Raimann A, Hojo H, Dong L, Roschger P, Kikani B, Wintergerst U, Fratzl-Zelman N, Jee YH, Haeusler G, et al. 2022. A neomorphic variant in SP7 alters sequence specificity and causes a high-turnover bone disorder. Nat Commun 13: 700.

Madsen BE, Browning SR. 2009. A groupwise association test for rare mutations using a weighted sum statistic. PLoS genetics 5: e1000384.

McCaw ZR, O’Dushlaine C, Somineni H, Bereket M, Klein C, Karaletsos T, Casale FP, Koller D, Soare TW. 2023. An allelic-series rare-variant association test for candidate-gene discovery. The American Journal of Human Genetics 110: 1330–1342.

McClintock B. 1944. The relation of homozygous deficiencies to mutations and allelic series in maize. Genetics 29: 478.

McLaren W, Gil L, Hunt SE, Riat HS, Ritchie GRS, Thormann A, Flicek P, Cunningham F. 2016. The ensembl variant effect predictor. Genome biology 17: 1–14.

Morales JL, Nocedal J. 2011. L-BFGS-B: Remark on Algorithm 778: L-BFGS-B, FORTRAN routines for large scale bound constrained optimization. ACM Transactions on Mathematical Software 38.

Musunuru K, Kathiresan S. 2019. Genetics of common, complex coronary artery disease. Cell 177: 132–145.

Neale B. 2018. UK Biobank GWAS results. Neale Lab. http://www.nealelab.is/uk-biobank/.

Nguyen E, Poli M, Durrant MG, Thomas AW, Kang B, Sullivan J, Ng MY, Lewis A, Patel A, Lou A, et al. 2024. Sequence modeling and design from molecular to genome scale with Evo. http://dx.doi.org/10.1101/2024.02.27.582234.

Ranganath R, Gerrish S, Blei D. 2014. Black box variational inference. In Artificial intelligence and statistics, pp. 814–822, PMLR.

Rentzsch P, Witten D, Cooper GM, Shendure J, Kircher M. 2019. CADD: predicting the deleteriousness of variants throughout the human genome. Nucleic acids research 47: D886–D894.

Rezende DJ, Mohamed S, Wierstra D. 2014. Stochastic backpropagation and approximate inference in deep generative models. In International conference on machine learning, pp. 1278–1286, PMLR.

Ritchie GRS, Flicek P. 2015. Functional Annotation of Rare Genetic Variants. In Assessing Rare Variation in Complex Traits: Design and Analysis of Genetic Studies (eds. E. Zeggini and A. Morris), Springer, New York (NY).

Rives A, Meier J, Sercu T, Goyal S, Lin Z, Liu J, Guo D, Ott M, Zitnick CL, Ma J, et al. 2021. Biological structure and function emerge from scaling unsupervised learning to 250 million protein sequences. Proc Natl Acad Sci U S A 118: e2016239118.

Sundaram L, Gao H, Padigepati SR, McRae JF, Li Y, Kosmicki JA, Fritzilas N, Hakenberg J, Dutta A, Shon J, et al. 2018. Predicting the clinical impact of human mutation with deep neural networks. Nature genetics 50: 1161–1170.

Wagner N, Çelik MH, Hölzlwimmer FR, Mertes C, Prokisch H, Yépez VA, Gagneur J. 2023. Aberrant splicing prediction across human tissues. Nature genetics 55: 861–870.

Wolf FA, Angerer P, Theis FJ. 2018. SCANPY: large-scale single-cell gene expression data analysis. Genome biology 19: 1–5.

Wu MC, Lee S, Cai T, Li Y, Boehnke M, Lin X. 2011. Rare-variant association testing for sequencing data with the sequence kernel association test. Am J Hum Genet 89: 82–93.

Yoshida CA, Komori H, Maruyama Z, Miyazaki T, Kawasaki K, Furuichi T, Fukuyama R, Mori M, Yamana K, Nakamura K, et al. 2012. SP7 inhibits osteoblast differentiation at a late stage in mice. PLoS One 7: e32364.

Zhou J, Troyanskaya OG. 2015. Predicting effects of noncoding variants with deep learning--based sequence model. Nature methods 12: 931–934.

Zhou W, Bi W, Zhao Z, Dey KK, Jagadeesh KA, Karczewski KJ, Daly MJ, Neale BM, Lee S. 2022. SAIGE-GENE+ improves the efficiency and accuracy of set-based rare variant association tests. Nature genetics 54: 1466–1469.

Zhu C, Byrd RH, Nocedal J. 1997. L-BFGS-B: Algorithm 778: L-BFGS-B, FORTRAN routines for large scale bound constrained optimization (1997). ACM Transactions on Mathematical Software 23: 550–560.

