## Supplementary information for "Bayesian Aggregation of Multiple Annotations Enhances Rare Variant Association Testing"

### Contents

**A1 Supplementary Information** **1**
A1.1 Related work . . . . . . . . . . . . . . . . . . . . . . . . . . . . . . . 1
A1.2 Simulation design . . . . . . . . . . . . . . . . . . . . . . . . . . . . . 2

**A2 Supplementary Figures** **3**

**A3 Supplementary Tables** **13**

### A1 Supplementary Information

#### A1.1 Related work

**Burden tests.** While our Bayesian RVAT framework is general, we here specialize it to perform burden tests. Burden tests aggregate the effects of rare variants into a gene burden score to improve statistical power. This score is then regressed against a trait of interest in a formal test [3, 13, 12, 18, 22, 19]. While traditional burden tests have been limited to few annotations such as minor allele frequency (MAF) and variant consequences [7, 13], BayesRVAT can integrate multiple annotations leveraging a Bayesian framework.

**Allelic series.** BayesRVAT can model gene-level effects as a function of multiple rare variants and their functional annotations, effectively capturing allelic series. The term "allelic series" refers to a collection of variants within a gene that exhibit a gradation of phenotypic effects based on their severity, suggesting a dose-response relationship between gene functionality and the resulting phenotype [21, 20]. As allelic series enable the assessment of the feasibility of pharmacological modulation [20, 5, 16], methods that can accurately capture these relationships are of significant interest. For example, COAST models allelic series by weighting variants based on the expected deleteriousness of few functional consequences [16]. Conversely, DeepRVAT uses a data-driven approach to learn a trait-gene-agnostic aggregation function from multiple annotations using neural networks [4]. In contrast to these methods, BayesRVAT can handle larger sets of variant annotations without enforcing a universally fixed scheme across genes and traits.

**Variance component models.** In contrast to burden tests, which assume a uniform effect direction across all variants, variance component approaches allow for both deleterious and protective effects by employing random effect models. The most widely used variance component test for rare variants is SKAT [27]. Given the complementary strengths of burden and variance component tests [1], omnibus tests that combine both, such as SKAT-O [11], have become increasingly popular[17, 14, 31]. In this work, we demonstrate that BayesRVAT integrates smoothly within omnibus test procedures, maintaining its power advantages over other integrated burden tests.

**Bayesian inference.** BayesRVAT performs Bayesian inference on parameters modeling variant effects as a function of multiple annotations. Given the intractability of exact posterior computation, we use black-box variational inference [23], which reformulates the inference problem as an optimization task, directly optimizing a variational distribution to approximate the true posterior using gradient-based methods [2, 24, 10, 6, 25]. While Bayesian methods for RVAT have been previously explored [29, 26, 15, 30], BayesRVAT is the first unified Bayesian framework that can incorporate multiple genetic architectures and variant annotations.

### A1.2 Simulation design

Briefly, for each random seed, we simulated the phenotype as follows:

- We randomly selected $C$ contributing continuous annotations;
- The gene burden was built using the additive model with saturation in Eq **??**, where the effects of pLoF and missense were sampled from their respective priors, and the effects of the $C$ selected contributing continuous annotations were sampled from $\mathcal{N}(4, 0.01)^*$;
- The burden was standardized to have a mean of 0 and standard deviation of 1 across individuals;
- The phenotype $\boldsymbol{y} \in \mathbb{R}^{N\times 1}$ was simulated as:

$$\boldsymbol{y} = \sqrt{v_g} \cdot \boldsymbol{b} + \sqrt{1 - v_g} \cdot \boldsymbol{\psi}, \quad \boldsymbol{\psi} \overset{\text{iid}}{\sim} \mathcal{N}(0, 1), \tag{A.1}$$

  where $\boldsymbol{b} \in \mathbb{R}^{N\times 1}$ is the standardized burden and $v_g$ represents the proportion of variance explained by the genetic component.

In this framework, we evaluated power of BayesRVAT and other burden tests varying $C$, $v_g$, and the sample size.

*This ensures that the selected contributing annotations have a measurable effect on the burden score, thus effectively controlling the number of contributing continuous annotations $C$.

### A2 Supplementary Figures

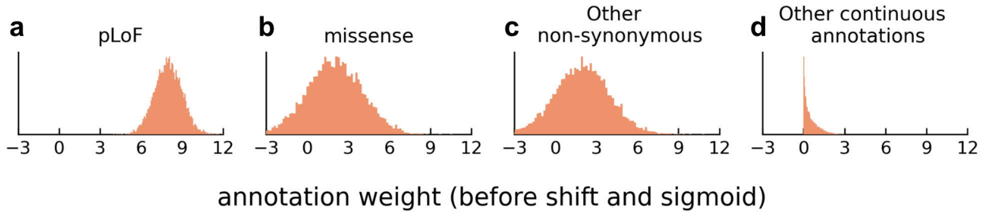


Figure A1: **Prior distributions for variant annotations used in BayesRVAT.** (**a**–**d**) Prior distributions over the raw annotation weights $\boldsymbol{\phi}$, before applying the shift $b_0$ and the sigmoid transformation in the aggregation function $g_{\boldsymbol{\phi}}(X, A) = \sigma(\boldsymbol{X}\boldsymbol{A}\boldsymbol{\phi} - b_0\mathbf{1}_{N\times 1})$. (**a**) pLoF variants are modeled with a normal prior (mean = 8, standard deviation = 1). (**b**) Missense variants are modeled with a normal prior (mean = 2, standard deviation = 2). (**c**) Other non-synonymous variants follow a normal prior (mean = 2, standard deviation = 2). (**d**) Continuous annotations, including regulatory and functional scores, are modeled with softplus-transformed Gaussian priors (mean = 0, standard deviation = 1, $\beta = 2$), ensuring positive contributions to the burden score.

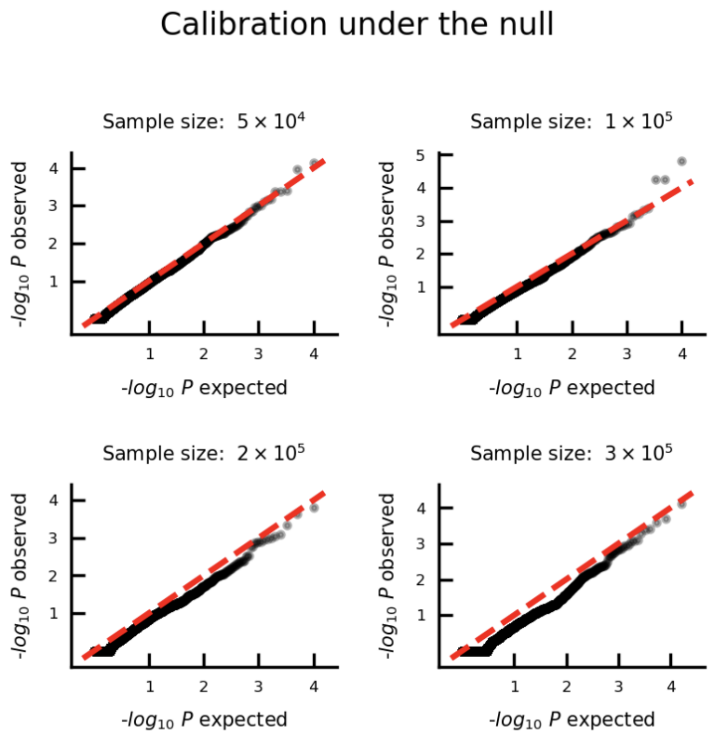


Figure A2: **Assessment of calibration of BayesRVAT in simulations varying sample size.** Shown are QQ plots of $P$ values from BayesRVAT under a null model with no genetic effects, using simulated phenotypes for 50,000, 100,000, 200,000, and 300,000 unrelated individuals from the UK Biobank (**Methods**).

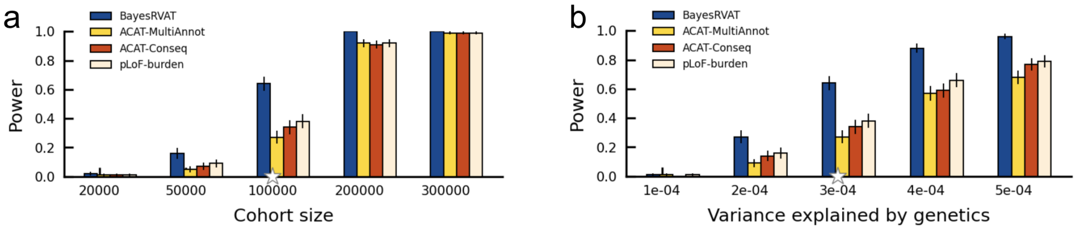


Figure A3: **Power assessment of BayesRVAT across different cohort sizes and variance explained by genetic effects.** (**a-b**) Statistical power of BayesRVAT compared to pLoF-burden, ACAT-Conseq, and ACAT-MultiAnnot across varying cohort sizes (**a**) and variance explained by genetic effects (**b**). Power is measured at the exome-wide significance threshold of $P < 2.5 \times 10^{-6}$, with results averaged over 100 random seeds for each setting.

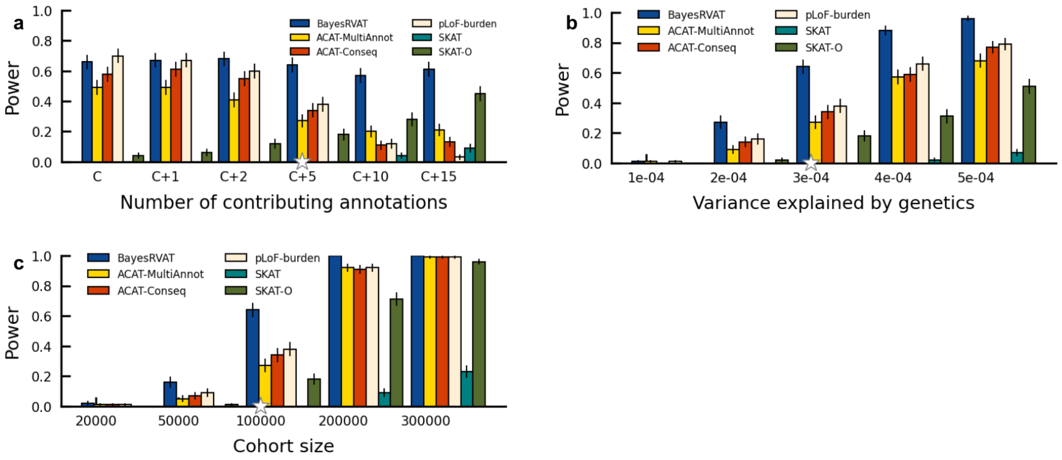


Figure A4: **Power assessment of BayesRVAT across different contributing annotations to the phenotype, cohort sizes and variance explained by genetic effects.** (**a-c**) Statistical power of BayesRVAT compared to pLoF-burden, ACAT-Conseq, ACAT-MultiAnnot, SKAT, and SKAT-O across varying number of contributing annotation to the phenotype (**a**) variance explained by genetics (**b**) and cohort size (**c**). Power is measured at the exome-wide significance threshold of $P < 2.5 \times 10^{-6}$, with results averaged over 100 random seeds for each setting.

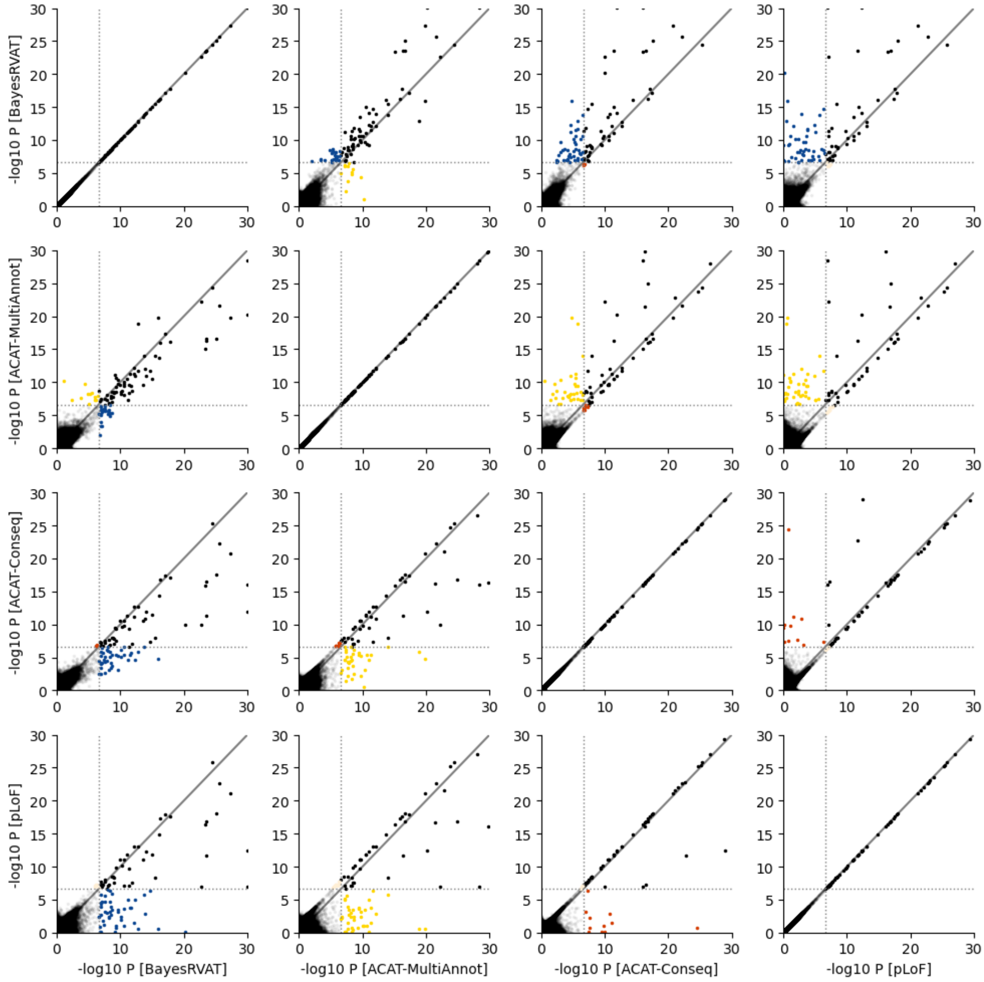


Figure A5: **Comparison of P-values from different burden test strategies in the blood trait analysis.** Each scatter plot compares the $-log_{10}$ P-values from two different methods, where points represent individual gene-trait pairs. Diagonal lines indicate equal P-values for the two methods being compared. Blue, yellow, red and pink points highlight gene-trait pairs uniquely identified by BayesRVAT, ACAT-MultiAnnot, and ACAT-Conseq, and pLoF, respectively, that are not detected by the other method.

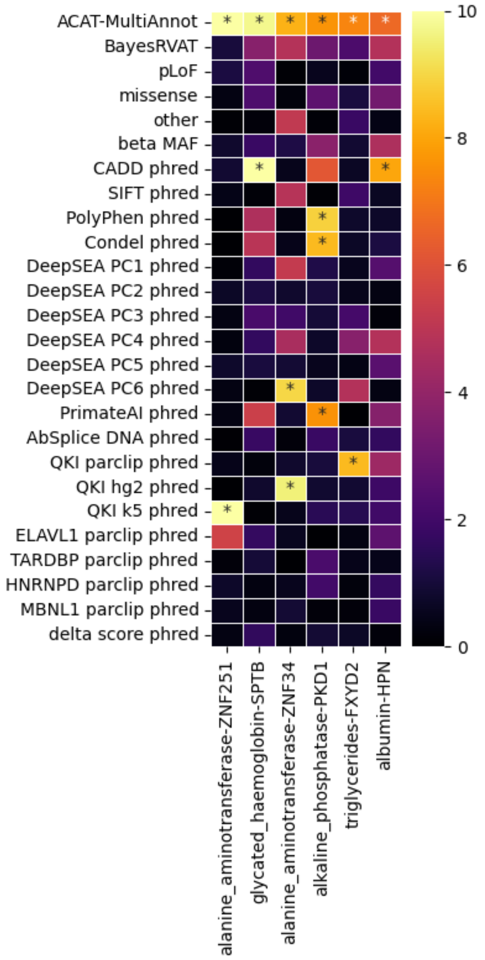


Figure A6: **Violations of the allelic series assumptions in BayesRVAT explain differences with ACAT-MultiAnnot.** The heatmap displays $-log_{10}$ P-values for gene-trait associations across individual annotations tested in ACAT-MultiAnnot, as well as the overall results for BayesRVAT and ACAT-MultiAnnot. Each column corresponds to a gene-trait pair identified by ACAT-MultiAnnot but missed by BayesRVAT, and each row represents a different annotation or burden test. In all these cases, the loss of power for BayesRVAT can be explained by the violation of the allelic series assumption encoded in its prior—namely, that pLoF variants have stronger effects than other annotations. In these gene-trait pairs, certain annotations (e.g., CADD, PolyPhen, Condel) exhibit much stronger effects than pLoF, driving the associations in ACAT-MultiAnnot. Stars (*) denote annotations with P-values that pass the Bonferroni-adjusted significance threshold of $\alpha < 0.05$.

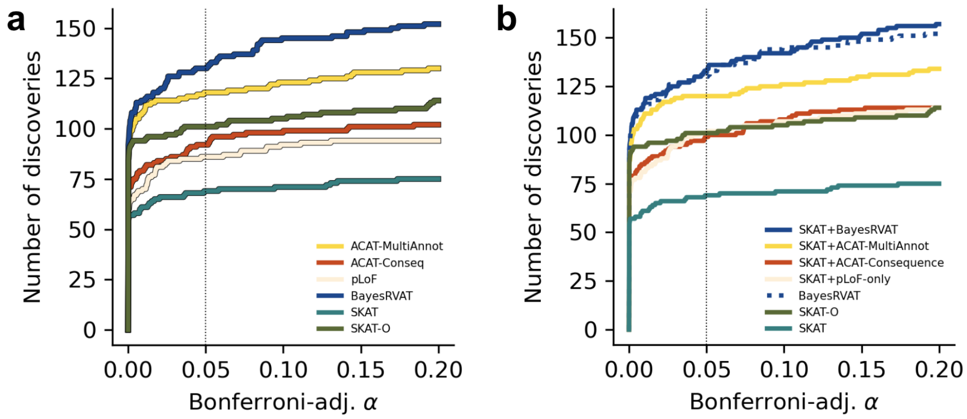


Figure A7: **Analysis of blood biomarkers in the UK Biobank including SKAT and SKAT-O.** (**a**) BayesRVAT outperform both burden (pLoF-Burden, ACAT-Conseq, and ACAT-MultiAnnot), variance components (SKAT) and optimal (SKAT-O) tests in number of discoveries at varying Bonferroni-adjusted significance thresholds $\alpha$. (**b**) BayesRVAT shows a superior number of discoveries even when other methods are integrated with SKAT results.

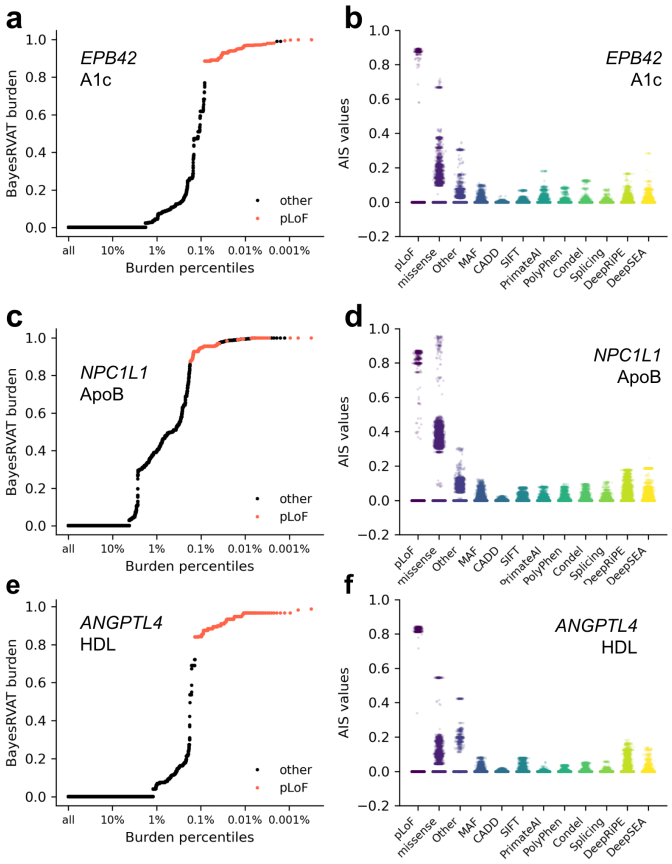


Figure A8: **Burden score and annotation importance scores (AIS) for additional blood biomarker associations detected by BayesRVAT.** (**a**) Burden scores learned by BayesRVAT for *EPB42* and HbA1c across burden percentiles, with individuals carrying pLoF mutations highlighted in red. This association aligns with recent findings showing a strong link between rare variants in *EPB42* and A1c levels [9]. (**b**) Annotation importance scores (AIS) for the association between *EPB42* and HbA1c, showing contributions from annotations such as missense, SIFT, and DeepSEA. (**c**) Burden scores for *NPC1L1* and apolipoprotein B (ApoB), with pLoF variants highlighted in red. *NPC1L1* is crucial for cholesterol absorption in the intestine and liver, and variants in this gene have been linked to altered ApoB levels, influencing lipid metabolism and cardiovascular disease risk [8]. (**d**) AIS for the association between *NPC1L1* and ApoB, highlighting the roles of missense and regulatory annotations. (**e**) Burden scores for *ANGPTL4* and HDL cholesterol, with pLoF variants highlighted in red. *ANGPTL4* is a key regulator of lipid metabolism, inhibiting lipoprotein lipase, which affects triglyceride breakdown and HDL cholesterol levels, with certain variants associated with increased HDL and cardiovascular protection[28]. (**f**) AIS for the association between *ANGPTL4* and HDL cholesterol, showing contribution from annotations such as missense, other non-synonymous, DeepRiPE, and DeepSEA.

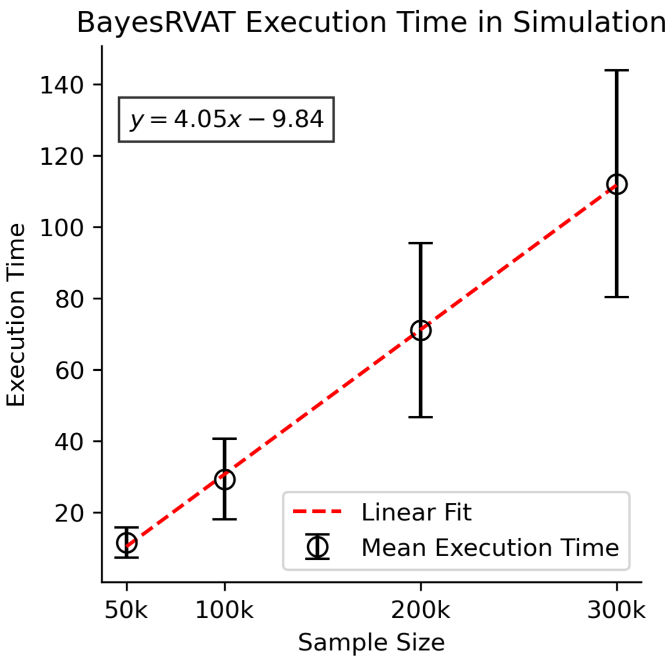


Figure A9: **BayesRVAT execution time for a single gene as a function of sample size.** The black circles represent the mean execution time for each sample size (50k, 100k, 200k, and 300k), with error bars indicating the standard deviation. A linear regression (red dashed line) is fitted to the data, illustrating the linear relationship between execution time and sample size.

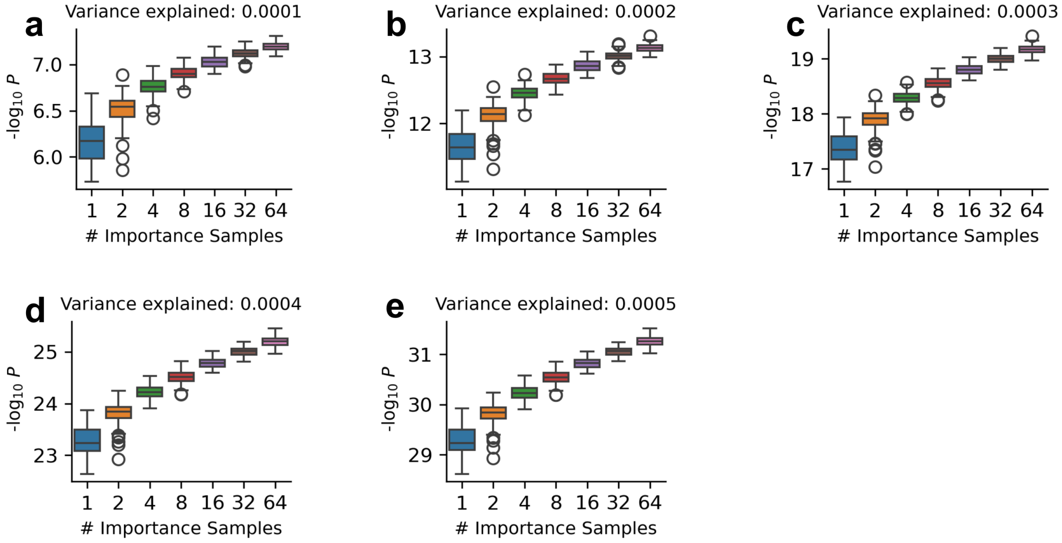


Figure A10: **Impact of importance sample size on the accuracy of $P$ value estimation.** Shown are distributions of $-\log_{10} P$ across 100 simulated phenotypes, evaluated under five settings with increasing levels of genetic variance explained. For each setting, we approximated the outer expectation in the IW-ELBO using 64 Monte Carlo samples, while varying the number of importance samples $K$ from 1 to 64. Increasing $K$ leads to higher and more stable test statistics, reflecting improved approximation of the (intractable) marginal likelihood under the alternative hypothesis and, consequently, more accurate $P$ values. In our experiments, we selected $K = 16$ as a practical trade-off between accuracy and computational cost.

### A3 Supplementary Tables

| Gene | Trait | $P_{\text{pLoF}}$ | $P_{\text{ACAT-Conseq}}$ | $P_{\text{ACAT-MultiAnnot}}$ | $P_{\text{BayesRVAT}}$ | $\beta_{\text{pLoF}}$ | $\beta_{\text{BayesRVAT}}$ |
|---|---|---|---|---|---|---|---|
| MIP | Cataract | $4.33 \times 10^{-13}$ | $1.30 \times 10^{-12}$ | $8.84 \times 10^{-28}$ | $7.54 \times 10^{-28}$ | 2.47 | 3.01 |
| GCK | T2D | $2.81 \times 10^{-16}$ | $8.88 \times 10^{-16}$ | $3.97 \times 10^{-22}$ | $3.60 \times 10^{-25}$ | 3.55 | 3.51 |
| LDLR | CAD | $1.12 \times 10^{-15}$ | $3.44 \times 10^{-15}$ | $2.84 \times 10^{-14}$ | $1.06 \times 10^{-21}$ | 2.18 | 2.05 |
| HNF1A | T2D | $1.62 \times 10^{-7}$ | $4.87 \times 10^{-7}$ | $1.92 \times 10^{-8}$ | $2.93 \times 10^{-14}$ | 1.73 | 3.32 |
| GUCY1A1 | HT | $2.47 \times 10^{-7}$ | $7.41 \times 10^{-7}$ | $3.62 \times 10^{-8}$ | $2.72 \times 10^{-12}$ | 0.92 | 1.01 |
| GIGYF1 | T2D | $4.82 \times 10^{-12}$ | $1.44 \times 10^{-11}$ | $1.20 \times 10^{-10}$ | $6.98 \times 10^{-12}$ | 1.71 | 1.90 |
| PRPH2 | AMD | $6.33 \times 10^{-7}$ | $1.90 \times 10^{-6}$ | $1.45 \times 10^{-6}$ | $8.52 \times 10^{-12}$ | 1.43 | 2.65 |
| FES | HT | $1.18 \times 10^{-3}$ | $1.41 \times 10^{-4}$ | $5.53 \times 10^{-8}$ | $2.37 \times 10^{-9}$ | 0.70 | 0.66 |
| PKD1 | HT | $2.25 \times 10^{-6}$ | $6.75 \times 10^{-6}$ | $5.63 \times 10^{-5}$ | $1.33 \times 10^{-7}$ | 1.15 | 2.44 |

Table A1: **Significant gene-trait associations from the analysis of eight disease traits.** Shown are P values for pLoF-only ($P_{\text{pLoF}}$), ACAT-Consequence ($P_{\text{ACAT-Conseq}}$), ACAT-MultiAnnot ($P_{\text{ACAT-MultiAnnot}}$), and BayesRVAT ($P_{\text{BayesRVAT}}$), and effect size estimates ($\beta$) for pLoF and BayesRVAT burdens.

### References


[1] Saonli Basu and Wei Pan. Comparison of statistical tests for disease association with rare variants. Genetic epidemiology, 35(7):606–619, 2011.

[2] David M Blei, Alp Kucukelbir, and Jon D McAuliffe. Variational inference: A review for statisticians. Journal of the American statistical Association, 112(518): 859–877, 2017.

[3] Elizabeth T Cirulli, Simon White, Robert W Read, Gai Elhanan, William J Metcalf, Francisco Tanudjaja, Donna M Fath, Efren Sandoval, Magnus Isaksson, Karen A Schlauch, et al. Genome-wide rare variant analysis for thousands of phenotypes in over 70,000 exomes from two cohorts. Nature communications, 11 (1):542, 2020.

[4] Brian Clarke, Eva Holtkamp, Hakime Öztürk, Marcel Mück, Magnus Wahlberg, Kayla Meyer, Felix Munzlinger, Felix Brechtmann, Florian R Hölzlwimmer, Jonas Lindner, et al. Integration of variant annotations using deep set networks boosts rare variant association testing. Nature Genetics, pages 1–10, 2024.

[5] Calliope A Dendrou, Adrian Cortes, Lydia Shipman, Hayley G Evans, Kathrine E Attfield, Luke Jostins, Thomas Barber, Gurman Kaur, Subita Balaram Kuttikkatte, Oliver A Leach, et al. Resolving tyk2 locus genotype-to-phenotype differences in autoimmunity. Science translational medicine, 8(363):363ra149–363ra149, 2016.

[6] Jan P Engelmann, Alessandro Palma, Jakub M Tomczak, Fabian Theis, and Francesco Paolo Casale. Mixed models with multiple instance learning. In International Conference on Artificial Intelligence and Statistics, pages 3664–3672. PMLR, 2024.

[7] Fang Han and Wei Pan. A data-adaptive sum test for disease association with multiple common or rare variants. Human heredity, 70(1):42–54, 2010.

[8] Lin Jia, Jenna L Betters, and Liqing Yu. Niemann-pick c1-like 1 (npc1l1) protein in intestinal and hepatic cholesterol transport. Annual review of physiology, 73 (1):239–259, 2011.

[9] Young Jin Kim, Sanghoon Moon, Mi Yeong Hwang, Sohee Han, Hye-Mi Jang, Jinhwa Kong, Dong Mun Shin, Kyungheon Yoon, Sung Min Kim, Jong-Eun Lee, et al. The contribution of common and rare genetic variants to variation in metabolic traits in 288,137 east asians. Nature communications, 13(1):6642, 2022.

[10] Diederik P Kingma. Auto-encoding variational bayes. arXiv preprint arXiv:1312.6114, 2013.

[11] Seunggeun Lee, Michael C Wu, and Xihong Lin. Optimal tests for rare variant effects in sequencing association studies. Biostatistics, 13(4):762–775, 2012.

[12] Seunggeung Lee, Gonçalo R Abecasis, Michael Boehnke, and Xihong Lin. Rare-variant association analysis: study designs and statistical tests. The American Journal of Human Genetics, 95(1):5–23, 2014.

[13] Bingshan Li and Suzanne M Leal. Methods for detecting associations with rare variants for common diseases: application to analysis of sequence data. The American Journal of Human Genetics, 83(3):311–321, 2008.

[14] Xihao Li, Zilin Li, Hufeng Zhou, Sheila M Gaynor, Yaowu Liu, Han Chen, Ryan Sun, Rounak Dey, Donna K Arnett, Stella Aslibekyan, et al. Dynamic incorporation of multiple in silico functional annotations empowers rare variant association analysis of large whole-genome sequencing studies at scale. Nature genetics, 52 (9):969–983, 2020.

[15] Benjamin A Logsdon, James Y Dai, Paul L Auer, Jill M Johnsen, Santhi K Ganesh, Nicholas L Smith, James G Wilson, Russell P Tracy, Leslie A Lange, Shuo Jiao, et al. A variational bayes discrete mixture test for rare variant association. Genetic epidemiology, 38(1):21–30, 2014.

[16] Zachary R McCaw, Colm O'Dushlaine, Hari Somineni, Michael Bereket, Christoph Klein, Theofanis Karaletsos, Francesco Paolo Casale, Daphne Koller, and Thomas W Soare. An allelic-series rare-variant association test for candidate-gene discovery. The American Journal of Human Genetics, 110(8):1330–1342, 2023.

[17] Remo Monti, Pia Rautenstrauch, Mahsa Ghanbari, Alva Rani James, Matthias Kirchler, Uwe Ohler, Stefan Konigorski, and Christoph Lippert. Identifying interpretable gene-biomarker associations with functionally informed kernel-based tests in 190,000 exomes. Nature communications, 13(1):5332, 2022.

[18] Stephan Morgenthaler and William G Thilly. A strategy to discover genes that carry multi-allelic or mono-allelic risk for common diseases: a cohort allelic sums test (cast). Mutation Research/Fundamental and Molecular Mechanisms of Mutagenesis, 615(1-2):28–56, 2007.

[19] Andrew P Morris and Eleftheria Zeggini. An evaluation of statistical approaches to rare variant analysis in genetic association studies. Genetic epidemiology, 34 (2):188–193, 2010.

[20] Kiran Musunuru and Sekar Kathiresan. Genetics of common, complex coronary artery disease. Cell, 177(1):132–145, 2019.

[21] Robert M Plenge, Edward M Scolnick, and David Altshuler. Validating therapeutic targets through human genetics. Nature reviews Drug discovery, 12(8): 581–594, 2013.

[22] Alkes L. Price, Gregory V. Kryukov, Paul I.W. de Bakker, Shaun M. Purcell, Jeff Staples, Lee-Jen Wei, and Shamil R. Sunyaev. Pooled association tests for rare variants in exon-resequencing studies. The American Journal of Human Genetics, 86(6):832–838, 2010. ISSN 0002-9297. doi: https://doi.org/10.1016/j.ajhg.2010.04.005. URL https://www.sciencedirect.com/science/article/pii/S0002929710002077.

[23] Rajesh Ranganath, Sean Gerrish, and David Blei. Black box variational inference. In Artificial intelligence and statistics, pages 814–822. PMLR, 2014.

[24] Danilo Jimenez Rezende, Shakir Mohamed, and Daan Wierstra. Stochastic backpropagation and approximate inference in deep generative models. In International conference on machine learning, pages 1278–1286. PMLR, 2014.

[25] Valentine Svensson, Adam Gayoso, Nir Yosef, and Lior Pachter. Interpretable factor models of single-cell rna-seq via variational autoencoders. Bioinformatics, 36(11):3418–3421, 2020.

[26] Guhan Ram Venkataraman, Christopher DeBoever, Yosuke Tanigawa, Matthew Aguirre, Alexander G Ioannidis, Hakhamanesh Mostafavi, Chris CA Spencer, Timothy Poterba, Carlos D Bustamante, Mark J Daly, et al. Bayesian model comparison for rare-variant association studies. The American Journal of Human Genetics, 108(12):2354–2367, 2021.

[27] Michael C Wu, Seunggeun Lee, Tianxi Cai, Yun Li, Michael Boehnke, and Xihong Lin. Rare-variant association testing for sequencing data with the sequence kernel association test. The American Journal of Human Genetics, 89(1):82–93, 2011.

[28] Long-Yan Yang, Cai-Guo Yu, Xu-Hong Wang, Sha-Sha Yuan, Li-Jie Zhang, Jia-Nan Lang, Dong Zhao, and Ying-Mei Feng. Angiopoietin-like protein 4 is a high-density lipoprotein (hdl) component for hdl metabolism and function in nondiabetic participants and type-2 diabetic patients. Journal of the American Heart Association, 6(6):e005973, 2017.

[29] Yi Yang, Saonli Basu, and Lin Zhang. A bayesian hierarchically structured prior for rare-variant association testing. Genetic epidemiology, 45(4):413–424, 2021.

[30] Nengjun Yi and Degui Zhi. Bayesian analysis of rare variants in genetic association studies. Genetic epidemiology, 35(1):57–69, 2011.

[31] Wei Zhou, Wenjian Bi, Zhangchen Zhao, Kushal K Dey, Karthik A Jagadeesh, Konrad J Karczewski, Mark J Daly, Benjamin M Neale, and Seunggeun Lee. Saige-gene+ improves the efficiency and accuracy of set-based rare variant association tests. Nature genetics, 54(10):1466–1469, 2022.
